# Optimizing the Fabrication of Shape-Defined Microparticles for Sustained Drug Delivery: the ‘Less is More’ paradigm

**DOI:** 10.1101/2024.09.10.612107

**Authors:** Denise Murgia, Bianca Martins Estevão, Corinne Portioli, Roberto Palomba, Paolo Decuzzi

**Author notes:** Both authors contributed equally (Denise Murgia, Bianca Martins Estevão). Corresponding author: Paolo Decuzzi, PhD –.

## Abstract

Polymeric microparticles find extensive use in several pharmaceutical applications. Our group has developed poly(lactic-co-glycolic acid) (PLGA) microPLates (μPL) featuring a square base of 20×20 μm and a height of 10 μm, for the controlled and sustained delivery of a range of therapeutic payloads, including anti-inflammatory and anti-cancer drugs, small molecules for neurodevelopmental disorders, and siRNA for osteoarthritis. In this study, the morphological and pharmacological properties of PLGA-μPL were optimized by introducing new steps in the original fabrication protocol and systematically varying the polymer content. Vacuum suction was used to control solvent removal, and two different ‘cleaning’ steps were tested, resulting in six different μPL configurations with a PLGA content ranging from 2 to 10 mg. Electron and optical microscopy analyses confirmed the well-defined square shape of μPL, with a central concavity depending on the PLGA content. Fabrication yielding ranged between 10% and 70%, while encapsulation efficiencies reached approximately 15% using curcumin (CURC) as a model drug. The kinetics of CURC release was analyzed using the semi-empirical model of Korsmeyer-Peppas, suggesting either a Fickian diffusion or anomalous transport mechanisms based on the PLGA amounts. Complementary techniques were used to assess morphological alterations and mass loss, evaluating the degradation μPL over time in water and physiological solutions. Unexpectedly, μPL configurations with lower PLGA contents exhibited higher fabrication yielding, drug encapsulation, and slower drug release. The optimized fabrication approach offers greater flexibility to tailor the degradation and pharmacological properties of μPL for various therapeutic applications.

## Introduction

There is a growing demand for novel platforms in the biomedical field, particularly in drug delivery, capable of improving therapeutic efficacy, accelerating recovery, and improving quality of life^1,2^. Polymeric nano/micro drug delivery systems offer precise control over release kinetics, tunability, and biocompatibility while preserving sensitive compounds from degradation^3,4^. Thus, the use of various polymers to design platforms with tailored shapes, sizes, and compositions has been intensely explored^5–9^.

Poly(lactic-co-glycolic acid) (PLGA) is a remarkable biodegradable and biocompatible copolymer derived from lactic acid (LA) and glycolic acid (GA). With its tunable drug release properties, it is employed to design versatile systems for delivering a wide range of therapeutics^10,11^. By adjusting the ratio of LA to GA, its composition can be modulated, affecting its crystallinity, hydrophobicity, and glass transition temperature. These manipulations influence the degradation rate and drug release kinetics, allowing customization for several therapeutic needs while ensuring minimal toxicity and immunogenicity^12^.

The concentration of polymers plays a crucial role in the development of systems for sustained drug delivery, as it influences the release kinetics and provides a means to tailor the duration and rate of the expected therapeutic effects^3,13–15^. Additionally, particle geometry significantly impacts drug release as it changes the volume-to-surface ratios and particle exposure to the surrounding environment, thus providing more tools to optimize the delivery of therapeutic agents^5,16^. Thus, the interplay between polymer concentrations and physical properties of drug delivery systems offers valuable strategies to optimize sustained release and design versatile platforms that meet specific application requirements and improve patient compliance^6^.

Our team has recently developed a microparticle-based drug delivery system consisting of shape-defined PLGA microparticles, called microPLates (μPL)^16–19^. These microcarriers have a unique geometrical structure with a square base of 20×20 μm and a height of 10 μm, and they have been used to deliver several therapeutic payloads, including anti-inflammatory and anti-cancer drugs, small molecules for neurodevelopmental disorders, and RNA-loaded nanoparticles for osteoarthritis management^16–20^. These μPL exhibit unique physicochemical attributes driven by their fixed geometry, surface characteristics, and mechanical features^17^. Unlike more conventional polymeric microspheres, these features can be tailored simultaneously and independently during a top-down microfabrication process, achieving uniformity and reproducibility, and holding the potential to confer specific advantages tailored to the intended application^16,17,20^.

In this study, an optimization of the μPL fabrication method using an under-vacuum approach and ad-hoc ‘cleaning’ steps is presented. The concentration of PLGA was systematically varied between 2 and 10 mg, while maintaining a constant input of the model drug, curcumin (CURC), chosen for its intrinsic fluorescence and well-known therapeutic properties^21,22^. Different μPL configurations were obtained and thoroughly characterized with respect to their physico-chemical and morphological properties, degradation behavior, drug loading and release profiles.

## Materials and methods

### Materials

Polydimethylsiloxane (PDMS) (Sylgard 184) elastomer was acquired from Dow Corning (Midland, Michigan, USA). Poly (vinyl alcohol) (PVA, MW 31,000 − 50,000), poly(D,L-lactide-co-glycolide), (PLGA, lactide:glycolide 50:50, MW 38,000 − 54,000), acetonitrile (ACN) and trifluoroacetic acid (TFA) were purchased from Sigma-Aldrich (Saint Louis, Missouri, USA). Polyphosphate Saline Buffer (PBS) was purchased from Gibco (Invitrogen Corporation, Giuliano Milanese, Milan, Italy). Curcumin (CURC) was obtained from Alfa Aesar (Haverhill, Massachusetts, USA). All reagents and solvents were used as received.

### Microfabrication of microPlates (μPL)

PLGA µPL were synthetized employing a multistep top-down fabrication approach, partially described in previous works of our group ^16–19^ and based on sequential replica molding steps, schematically reported in **Figures 1A** and **1B**. Initially, a silicon master template was fabricated using a direct laser writing technique (DLW), realizing well-defined geometries based on arrays of square wells. In the current configuration, the square wells had an edge length of 20 μm and a depth of 10 μm, separated by a 5 μm gap. Then, a PDMS replica was obtained by covering the silicon master template with a mixture of PDMS and a curing agent (10:1, v/v), removing the bubbles generated during the mixing step via a vacuum chamber, and baking in oven at ∼ 60 °C for 8 h to promote polymerization. Afterwards, the PDMS template was carefully peeled off the silicon master and used to obtain sacrificial PVA templates. The latter was created by pouring a PVA solution (5% w/v in deionized (DI) water) on the PDMS patterned surface. The sample was then dried at ∼ 60°C to obtain a PVA film that was finally peeled off the PDMS template, eventually presenting the same array of wells as the original silicon master mold. In the last fabrication step, the wells of the PVA template were filled by carefully spreading the PLGA and CURC polymeric paste with the help of a pipette tip. The paste was obtained by dissolving in ACN 0.375 mg of CURC and different amounts of polymer (2 mg, 5 mg, 7.5 mg, 10 mg of PLGA resulting in CURC-μPL-2, CURC-μPL-5, CURC-μPL-7.5 and CURC-μPL-10 configurations, respectively). Solvent evaporation was then controlled by leaving the samples in a vacuum-chamber for 10 min. Then, the loaded PVA template was dissolved in DI water at room temperature in an ultrasonic bath or, in the case of the μPL-10 configuration, it was either wiped with an ACN-soaked tissue (CURC-μPL-10-T configuration) or an ACN-soaked blade (CURC-μPL-10-B configuration). For all the configurations, the purification process then involved the use of polycarbonate membrane filters (40 μm pore size) to remove residual pieces of PVA. Finally, μPL were recovered after two sequential centrifugation steps (5,000 rpm for 5 min) and stored at 4 °C for further use. Within this process, six different CURC-μPL configurations were fabricated and characterized. μPL with different PLGA amounts (2 mg, 5 mg, 7.5 mg, 10 mg) were also prepared using the traditional and previously described method^16–19^, to compare their features with the presented μPL.

**Figure 1.**
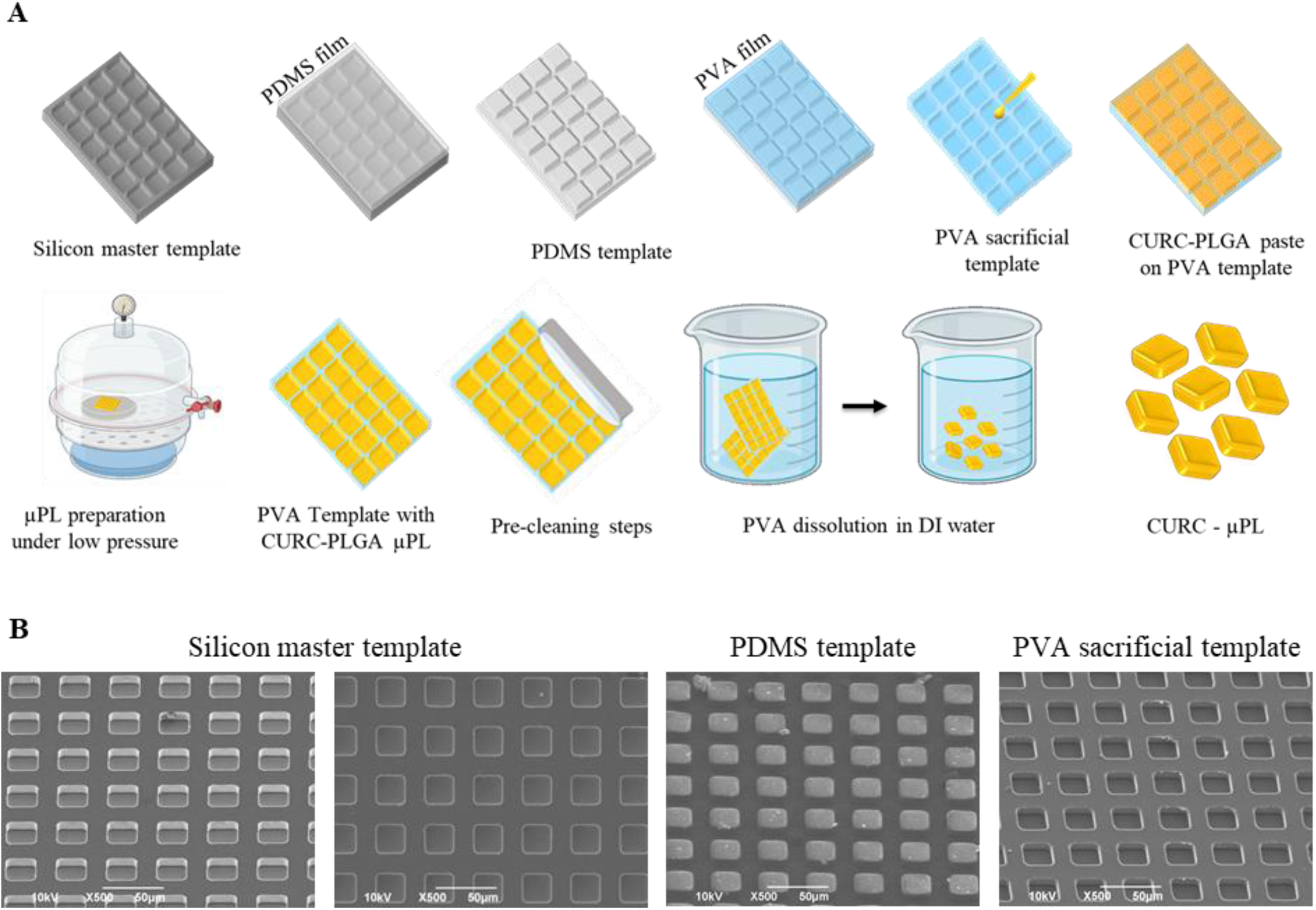
Top-down fabrication process for PLGA microPLates (μPL). **A.** Schematic representation of the steps needed for the PLGA μPL fabrication. From top left to bottom right: a ‘silicon master template’ (dark gray) is fabricated via direct laser writing; replicated into a ‘PDMS template’ (light gray); and transferred into a ‘PVA sacrificial template’ (light blue). A polymeric paste including CURC (0.375 mg) and PLGA (2 mg – 10 mg) (yellow) is carefully spread over the PVA template and rapidly placed in a vacuum chamber. The resulting PVA templates are directly dissolved in DI water, except for two μPL configurations which undergo ‘cleaning’ (ACN-soaked tissue; ACN-wet blade) before this step. The released μPL (yellow) are collected by centrifugation; **B.** Scanning electron microscopy (SEM) images of the templates. From left to right: a 45° tilted view; a frontal view of the ‘silicon master template’; a 45° tilted view of the ‘PDMS template’; and a 45° tilted view of the ‘PVA sacrificial template’.

### Physical-chemical characterization of microPLates (μPL)

The morphological and physicochemical properties of μPL were evaluated using different techniques. μPL size and shape were documented via scanning electron microscopy (SEM, Elios Nanolab 650, FE), while particle morphology and the homogeneous CURC distribution were assessed via fluorescent microscopy (Nikon A1, Dexter, MI and Leica DM6000 B, Leica Microsystems, Wetzlar, DE).

For SEM imaging, a drop of sample was spotted on a silicon template and uniformly sputtered with 10 nm of gold to increase the contrast and reduce sample damage. Samples were analyzed at an acceleration voltage of 10 keV. For fluorescent microscopy imaging, a drop of the sample was dotted on a microscope slide and covered by a cover glass. These microscopy techniques were used to characterize the morphology of each individual μPL, and Silicon, PDMS and PVA templates.

The number of μPL per PVA template and size distribution were derived via a Multisizer 4 COULTER particle counter (Beckman Coulter, CA). Briefly, μPL were resuspended in an electrolyte solution and analyzed following protocols described by the vendor. The fabrication yielding was calculated using the following formula:

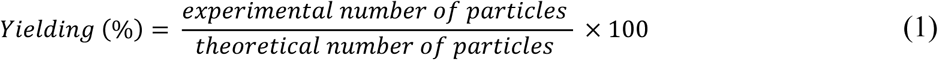

where the theoretical number of particles was calculated based on the design of the template, returning a value of 486,903 particles.

### Pharmaceutical properties of microPLates (μPL)

CURC loading, encapsulation efficiency (EE), and drug amounts per μPL were quantified as previously reported^16–19^. Briefly, CURC-μPL were lyophilized, dissolved in ACN/DI water (1:1, v/v), and analyzed by high-performance liquid chromatography (HPLC) (Agilent 1260 Infinity, Germany), equipped with a 100 μL sample loop injector and a C18 column (2.1 × 100 mm, 3.5 μm particle size, Agilent Eclipse Plus, USA) for chromatographic separation. For CURC elution, the mobile phase consisted of DI water acidified with TFA (0.1% v/v) and ACN acidified with TFA (0.1% v/v) pumped in isocratic flow rate of 0.3 mL/min at the ratio of 43:57 (v/v). The analysis was performed taking 430 nm as the detection wavelength and CURC was determined by interpolating a standard calibration curve in ACN/DI water (1:1, v/v) in the linear range of 0.1 − 50 µg/mL. No interferences were observed among CURC and PLGA and other components of the measured solution at the testing concentrations, and no change in drug elution peak was experienced in the presence of excipients. EE, loading, and amounts of CURC per μPL were defined using the following formula (2, 3, 4):

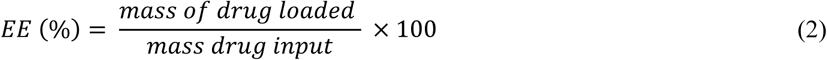

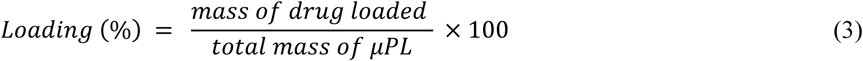

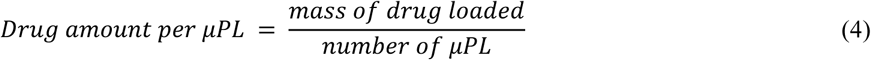

The CURC release profiles from each μPL configuration were determined up to one month. μPL were incubated in 500 μL of PBS solutions under magnetic stirring at 37 °C. At predetermined time points, samples in triplicate were collected and centrifuged (12,000 rpm for 5 min), discharging the supernatants. Pellets were dissolved in ACN/DI water (1:1, v/v) and analyzed by HPLC. The experimental release data were elaborated by GraphPad Prism 9 software (GraphPad Software, San Diego, CA, USA) and fitted to different models describing drug release kinetics^23,24^.

### Biodegradation of microPLates (μPL)

The μPL matrix degradation was assessed via SEM analysis and Multisizer Coulter counter. A 500 μL solution of μPL was incubated in PBS buffer (pH 7.4) and in DI water under mechanical stirring at 37 °C. At predetermined time points (1 day, 3 days, 7 days, 14 days and 28 days), samples were analyzed to monitor structural and morphological changes. The number of particles versus time was defined as Particle Number (%) via the following formula:

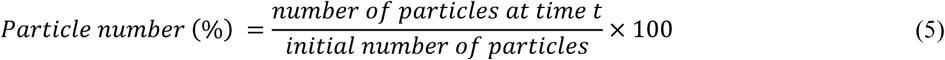

### Statistical Methods

Data were expressed as mean ± SD of at least three replicates (n = 3). For statistical analyses, Student’s t-test was applied with the minimum levels of significance being p < 0.05. One-way ANOVA followed by Dunnett’s multiple comparison tests were performed for all the pharmacological characterization using GraphPad Prism 9. Fittings of the drug release profiles were conducted using χ^2^ where a p-value of less than 0.05 was considered statistically significant.

## Results and Discussion

### Microfabrication and geometrical characterization of microPLates (μPL)

The original method for the fabrication of microPLates (μPL), described in previous studies by the authors^16–19^, was here extensively reviewed and improved to optimize various particle properties. The steps for μPL fabrication are sketched in **Figure 1A**. Briefly, the first step was the fabrication of the ‘silicon master template’ presenting a regular 2D array of microscopic wells which uniquely define the geometry of the μPL. This master template was then replicated into a PDMS intermediate template exhibiting a regular 2D array of pillars which was then replicated into a PVA sacrificial template. The PVA template presented a 2D array of square wells that perfectly matched the original geometry of the wells in the silicon master template. The well in the PVA template has an edge length of 20 μm and a height of 10 μm. This can be readily appreciated by reviewing the SEM images of **Figure 1B** showing, from left to right, a 45° inclined view and frontal views of the silicon master template with its wells; a 45° inclined view of the PDMS template with its pillars; and a 45° inclined view of the PVA sacrificial template with its wells. Then, a polymeric paste obtained by dissolving PLGA and CURC in ACN was evenly spread onto the PVA template, ensuring that the wells were carefully filled. A first modification to the original fabrication strategy consisted in rapidly positioning the so-loaded PVA template inside a vacuum chamber. This allowed the authors to control the evaporation of the organic solvent, without relying on the environmental conditions, resulting in a more uniform coverage of the template and accurate accumulation of the polymeric mixture within the wells. Then, to gently remove the backing layer of PLGA and CURC that inevitably forms over the PVA templates, different ‘cleaning’ steps were proposed. Specifically, the PVA template was gently wiped either with a blade or a tissue soaked in ACN or was not treated at all. Finally, the sacrificial PVA template was gradually dissolved in water, releasing the μPL that were collected via centrifugation.

This fabrication process was used to generate six distinct configurations of μPL by changing the amount of PLGA used per template and the ‘cleaning’ strategy. Specifically, μPL were obtained using 2 mg (CURC-μPL-2), 5 mg (CURC-μPL-5), 7.5 mg (CURC-μPL-7.5), and 10 mg (CURC-μPL-10) of PLGA.

### Physico-chemical characterization of microPLates (μPL)

The morphology of the μPL and the spatial distribution of the CURC molecules within the polymeric matrix were characterized using various microscopy techniques. The **Figures 2A** – C depict, respectively, scanning electron microscopy (SEM), confocal microscopy, and fluorescence microscopy images, of μPL synthesized using varying amounts of PLGA (from 2 mg to 10 mg) and employing different cleaning steps. All μPL configurations appeared as square prisms characterized by an edge length of approximately 20 μm and a height of about 10 μm, following the size of the wells in the silicon master and PVA sacrificial templates. SEM and fluorescent microscopy images revealed a well-formed structure with a concave top surface, which was especially evident in μPL-2 and μPL-10B. Note that, the μPL-2 were realized using the smallest amount of PLGA, likely causing an insufficient filling of the wells in the PVA sacrificial template; whereas the backing layer for the μPL-10B was removed – ‘cleaning’ step – via an ACN-wet blade, whose mechanical action could result in the partial removal of the paste from the wells of the PVA sacrificial template. The concavity of the top surface was less pronounced for the other four configurations where no treatment was applied to remove the backing layer (μPL-2, μPL-5, μPL-7.5, μPL-10) or an ACN-soaked tissue was used (μPL-10T). Therefore, the wiping of the PVA template with the blade (μPL-10B) appears to remove more materials than when using the tissue (μPL-10T). Also, the confocal microscopy images (**Figure 2B**), in agreement with the SEM (**Figure 2A**), showed more defined and less concave particles in the case of higher PLGA content. Moreover, the fluorescence images (**Figure 2C**) confirmed a uniform yellow-green coloration for the μPL, documenting the homogeneous distribution of CURC throughout the polymeric matrix. It should be here noted that the amount of CURC used to load the μPL was fixed for all configurations, and equal to 0.375 mg per template. Therefore, the lower green fluorescence associated with μPL-2 (**Figure 2C**, left) could be ascribed to quenching, due to CURC packing into a small volume.

**Figure 2.**
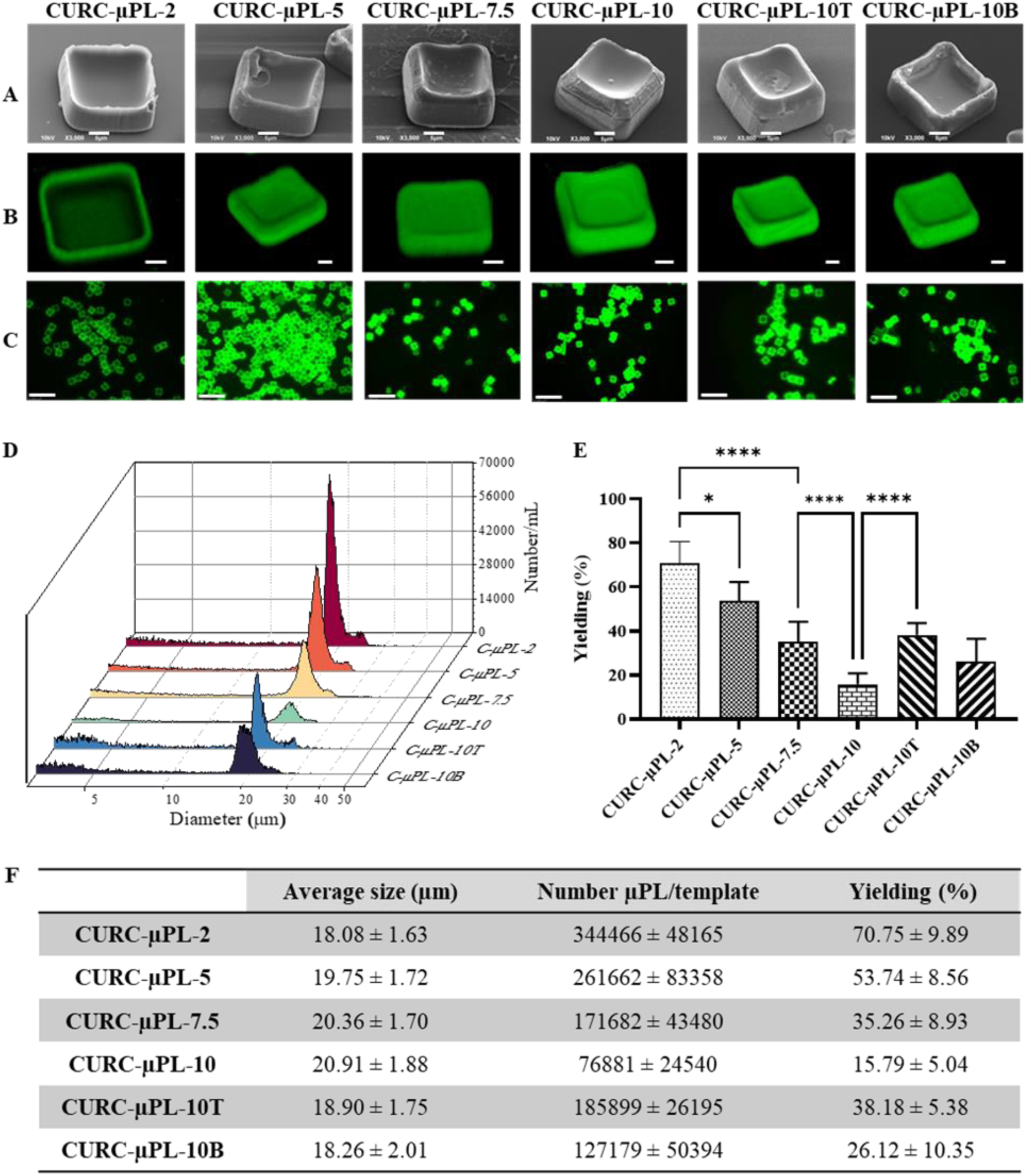
Morphological characterization of CURC-μPL. **A**. SEM images (45° tilted); **B.** 3D fluorescence confocal images (scale bar: 5 μm); **C**. Fluorescence microscopy images of the six different CURC-μPL configurations (scale bar: 100 μm); **D.** Multisizer Coulter Counter analyses of the six μPL configurations; **E.** Fabrication yielding; **F.** Summary of the CURC-μPL size, number per template, and fabrication yielding (Results are presented as average ± SD (n = 9). ****: p <0.0001).

To quantitatively assess the μPL morphology over the entire fabrication batch resulting from the dissolution of a PVA sacrificial template, a Multisizer Coulter counter system was employed. **Figure 2D** presents the size distribution profiles for aqueous solutions containing different CURC-μPL configurations. Importantly, data generated from a Multisizer Coulter counter are based on the assumption of spherical particles, therefore the peaks in **Figure 2D** correlate with an average size of μPL, while the value of the peak is instead related to the number of particles in solution. The μPL are well formed returning an average size close to their edge length (20 μm) regardless of the PLGA amounts and cleaning strategy. This was expected as the actual geometry of the μPL is defined by the original template, which is the same for all six μPL configurations. However, the fabrication yielding, calculated as the ratio between the number of fabricated μPL per template and the number of wells presented in the silicon master template (theoretical number of μPL per templates), differs significantly depending on the μPL configuration. The results are presented in **Figure 2E** and listed in the table in **Figure 2F**, together with values of the average particle size and the μPL number/template for each configuration. It results that the highest yielding is achieved for the CURC-μPL-2 returning a value of 70.75 ± 9.89%. As the PLGA content reduces, the yielding reduces becoming 53.76 ± 8.56% for the CURC-μPL-5, 38.18 ± 5.38% for the for the CURC-μPL-7.5, 35.26 ± 8.93% for the CURC-μPL-10T, 26.12 ± 10.35% for the CURC-μPL-10B and 15.79 ± 5.04% for the CURC-μPL-10. This data suggests that polymeric pastes with a lower PLGA content, and therefore less viscous, can be more accurately deposited within the wells of the PVA template limiting the formation of the backing layer, increasing the number of particles produced per template, and improving the particle morphological properties. This is evident also in **Figure 2D** with well-defined peaks in the Multisizer spectrum associated with the CURC-μPL-2. In contrast, the CURC-μPL-10 is associated with a modest yielding and the lowest peak in **Figure 2D**. One can observe that variations in polymer concentration influence the average size of microparticles. These results confirm that the mass of PLGA and the cleaning steps have a significant impact on the physicochemical properties of μPL.

### Pharmacological characterization of microPLates (μPL)

CURC was chosen as a model drug, as its intrinsic green-yellow fluorescence allows for quantification via conventional imaging techniques. Additionally, CURC also has multiple relevant pharmaceutical (anticancer, anti-inflammatory, and antioxidant) and nutraceutical properties as extensively documented in the open literature^25–28^. However, the therapeutic value is limited by its poor solubility in water, lack of stability, and low bioavailability, significant challenges that make it an ideal candidate for testing and developing advanced drug delivery systems^29^. The six μPL configurations were analyzed in terms of CURC encapsulation efficiency (EE – **Figure 3A**), loading (**Figure 3B**), and amount of CURC per μPL (**Figure 3C**). All the results were listed in a table (**Figure 3D**). The CURC input amount was fixed for all the μPL configurations being 0.375 mg. Following liquid chromatographic analyses, CURC-μPL-2 demonstrated the highest EE equal to 13.22 ± 1.57%. The efficiency in encapsulating CURC reduced with increased PLGA content becoming 10.66 ± 0.95%, 8.21 ± 1.90%, 3.99 ± 1.19%, 2.75 ± 0.58%, and 2.63 ± 0.70% for CURC-μPL-5, CURC-μPL-7.5, CURC-μPL-10, CURC-μPL-10T, and CURC-μPL-10B, respectively. A statistically significant difference was observed among the different PLGA configurations, confirming the importance of the polymer content. For the 10 mg PLGA case, no statistically significant difference was observed among the cleaning conditions (CURC-μPL-10: no cleaning; μPL-10T: with tissue, CURC-μPL-10B: with blade). A similar trend was also observed for CURC loading, where the maximum was again achieved for the CURC-μPL-2 and loading steadily decreased moving towards lower PLGA contents. It is here important to recall that the EE is defined as the ratio between the mass of drug actually entrapped into the μPL and the total input of drug per template, while loading is defined as the ratio between the mass of drug actually entrapped in the μPL and the total mass of the μPL (polymer and drug). While the total mass of CURC added per template is fixed, the μPL configurations with higher PLGA contents suffer from a thicker backing layer, resulting in higher drug losses and therefore lower encapsulation efficiencies. This is even more so for the loading. Interestingly, the mass of CURC loaded per particle, calculated as the ratio between the actual loaded amount of CURC per template divided by the number of μPL produced by that template, is almost constant (∼ 200 pg/μPL) for all the CURC-μPL configurations. Indeed, the lower PLGA μPL configurations (CURC-μPL-2) are characterized by high encapsulation and high fabrication yielding, whereas the higher PLGA μPL configurations (CURC-μPL-10) are characterized by low encapsulation and equally low fabrication yielding, so that the ratio between the two quantities stays approximately constant across all tested configurations. The CURC-μPL-10T do not follow the same trend as they are characterized by a low encapsulation and an intermediate yielding (∼ 40%).

**Figure 3.**
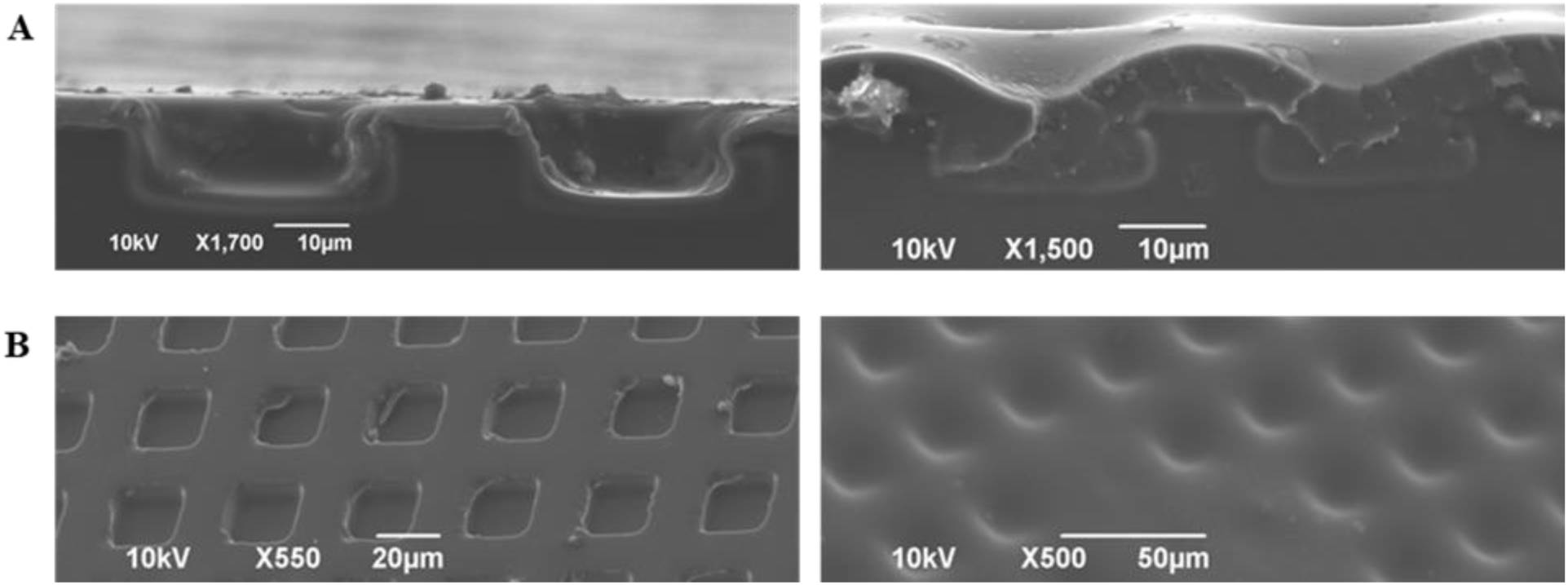
Effect of the polymer content on the formation of the ‘backing layer’. **A.** SEM images of the cross-section of PVA templates for CURC-µPL-2 (left) and CURC-µPL-10 (right), before cleaning; **B.** 45° tilted SEM images of the PVA templates for CURC-µPL-2 (left) and CURC-µPL-10 (right) before cleaning.

The six different CURC-μPL configurations were further investigated for their ability to release the encapsulated drug. CURC release was continuously monitored for up to 28 days (672 hours) under physiologically relevant conditions (PBS pH = 7.4, at 37 °C, with stirring) in a confined volume of 0.5 mL, mimicking the volume associated with an intra-tissue depot of μPL. The results of these release experiments are depicted in **Figure 3E** in terms of percentage of drug release as a function of time up to 28 days. The overall release rate was observed to increase with the PLGA content. CURC-µPL-2 exhibited the slowest and most sustained release, reaching a 61.75 ± 6.54% release fraction after 10 days followed by a plateau until the end of the test. For CURC-µPL-5, a 76.72 ± 2.86% release was documented at 10 days, while the CURC-μPL-7.5 and CURC-μPL-10 presented faster release rates, releasing over 80% of the loaded drug within the first 10 days. The µPL configurations that underwent the additional purification steps with tissue or blade soaking in ACN exhibited intermediate release rates. Drug release from PLGA-based particles is typically associated with two main mechanisms, namely molecular diffusion along the small pores of the polymeric matrix, dominating the process within the initial release phases, and degradation/erosion of the polymer fibers constituting the polymeric matrix, dominating the process during the latter release phase^30^. To gain more insights into how the different µPL configurations impact on CURC release, the commonly used semi-empirical model of Korsmeyer–Peppas was used to fit and interpret the experimental data^23,24^. The **Figure 3F** displays the fraction of drug released (M_t_/M_∞_) as a function of time and the fitting of the Korsmeyer-Peppas model, applicable only for the first 60% of the release curve table, while the **Figure 3G** lists the values for all the fitting parameters used in the analysis with the corresponding accuracy (R^2^)^31^. In the Korsmeyer–Peppas method, the exponent *n* plays a crucial role in characterizing drug release, as it accounts for both structural modifications and geometrical characteristics of the system. Approximating the μPL as a cylinder, for *n* = 0.45, the dominating mechanism of release is Fickian diffusion; for 0.45 < *n* < 0.89, non-Fickian or anomalous transport occurs where diffusion, degradation and swelling contribute to the process; for *n* > 0.89, another type of non-Fickian diffusion occurs, which was not observed in this study^23,24^. For low PLGA content, the CURC-μPL configurations were associated to *n* values close to 0.45 (0.43 ± 0.03 for CURC-μPL-2 and 0.40 ± 0.04 for CURC-μPL-5), confirming that Fickian diffusion dominates for these configurations. This process is characterized by solvent diffusion into the polymer matrix coupled with a low velocity of polymeric relaxation, promoting the diffusion of drug molecules from the core into the surrounding aqueous environment. This is indeed in agreement with the notion that lower PLGA contents would be associated to an overall lower hydrophobicity of the polymer matrix. Also, the lowest PLGA contents would limit local acidification (autocatalytic effect) and bulk degradation of the polymer matrix. All this would explain the lower release rates observed for the lower PLGA μPL configurations. Differently, with an increase in PLGA content, *n* became significantly larger than 0.45 (0.67 ± 0.03 and 0.58 ± 0.03 for CURC-μPL-7.5 and 10, respectively), suggesting anomalous transport as the drug release mechanisms, where polymer chain relaxation and Fickian diffusion are both involved^23,32^. Furthermore, where pre-purification methods were applied, the parameter *n* resulted equal to 0.45 ± 0.05 and 0.46 ± 0.05 for CURC-μPL-10T and CURC-μPL-10B respectively, shifting the release mechanism back to Fickian diffusion. These results confirm that the mass of PLGA and the cleaning steps have a significant impact on the pharmacological properties of CURC-μPL.

### Effect of the PLGA content and cleaning process on microPLates (μPL)

To gain deeper insights into how PLGA amounts, and cleaning steps could impact the morphology, physico-chemical, and pharmacological properties of μPL, SEM and confocal images were taken of the PVA templates loaded with the polymeric paste before cleaning. The two extreme configurations were considered, namely CURC-μPL-2 and CURC-μPL-10. SEM images were acquired for the template cross sections (**Figure 4A**) as well as a 45° tilted front (**Figure 4B**). The presence of a thicker backing layer for the CURC-μPL-10, resulting from a higher polymer amount, was evident in both images. This motivated the addition of a ‘cleaning step’ in the fabrication process, specifically for the high-polymer content configurations.

**Figure 4.**
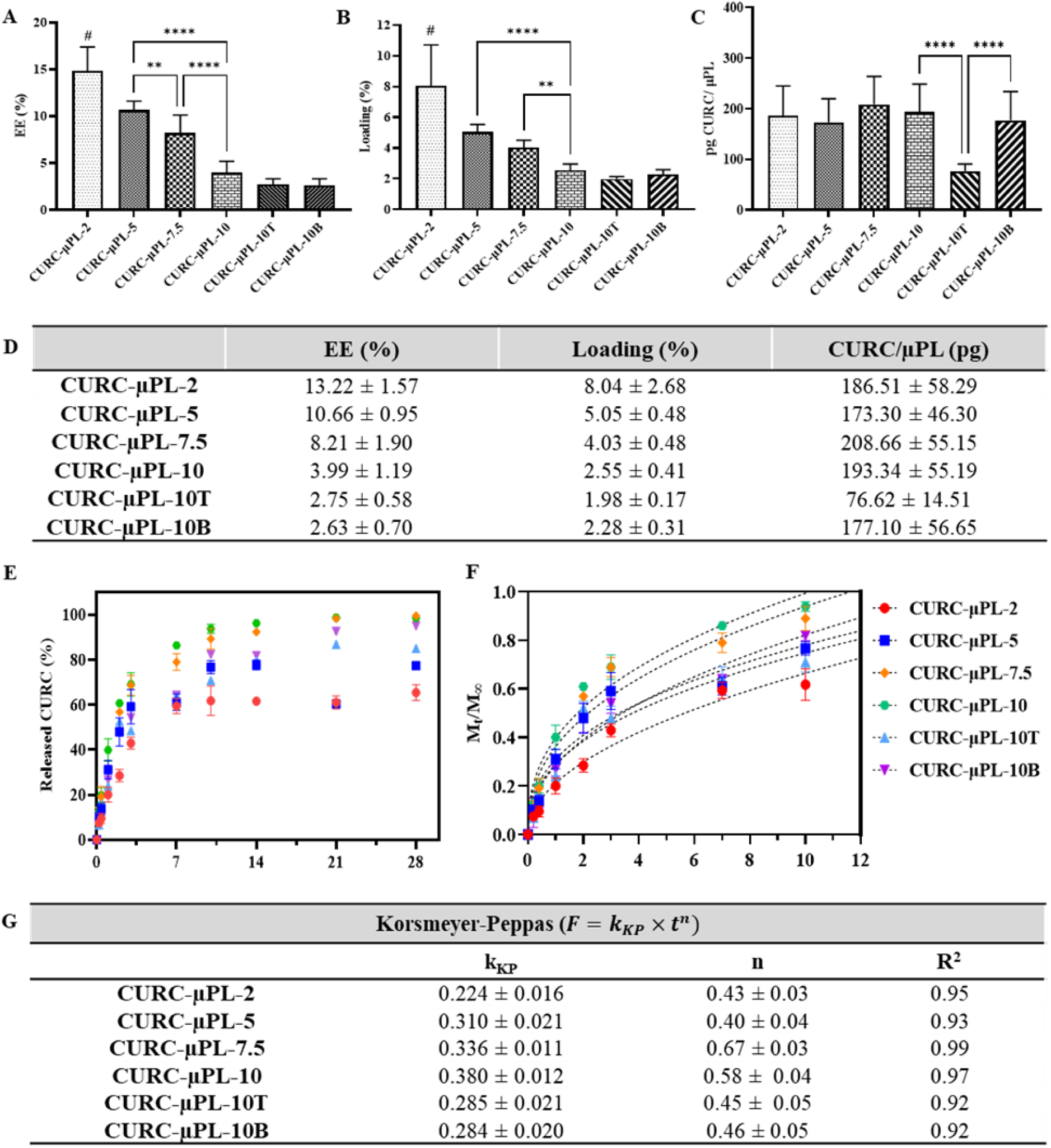
Pharmacological characterization of CURC-μPL. **A.** Encapsulation efficiency (EE); **B.** Loading; **C.** Amount of CURC per µPL (pg/µPL) for the six different configurations; **D**. Table summarizing the loading data; Results are presented as average ± SD (n = 9), ****: p <0.0001; # µPL-2 significant with all groups; **E.** *In vitro* CURC release in a confined volume (0.5 mL) of PBS buffer (pH = 7.4), for the six different configurations up to 28 days; **F.** Experimental data (colored dots) are fitted to Korsmeyer-Peppas (dashed lines) up to 60% of drug release; **G.** Values of the fitting parameters for the release kinetic model of Korsmeyer-Peppas. *Mt/M_∞_* equals the fraction of released drug within the time *t*; *k_KP_* is the release constant based on the structural characteristics of the drug-dosage form (here considered as cylinders); *n* is the diffusional exponent indicating the drug-release mechanisms. Results are presented as the average ± SD (n = 3).

Within this context, **Figure 5** presents a side-by-side comparison for the morphological and biopharmaceutic properties of μPL obtained with the newly proposed versus the original fabrication methods. SEM and confocal images confirmed for both μPL the unique squared shape, which is imparted by the master template regardless of the amount of polymer or cleaning process. However, the μPL obtained using the vacuum chamber and proper cleaning (**Figure 5A**) displayed a more regular and ‘filled’ shape, whereas the original μPL (**Figure 5B**) has a more pronounced concavity of the top surface and a larger percentage of empty, pierced, or incompletely filled structures. Also, the fluorescent microscopy showed a non-uniform distribution of the green signal of CURC among the particles, which was often not visible in the center (base) of the microparticles. Moreover, the data of **Figure 5C** confirms a higher fabrication yielding associated with the vacuum and cleaning steps especially for the lower PLGA amounts. At higher polymer contents, the differences between the two fabrication methods are not statistically significant. Even greater is the advantage in using the newly proposed approach when considering the EE and loading. **Figures 5D** and **5E**, respectively, document a four-fold increase and two-fold increase in EE and loading for the CURC-μPL-2 fabricated with the new method versus the original method. In general, EE and loading are higher for the new method across all tested configurations, as also observed in **Figure 5F** for the mass of CURC per particle.

**Figure 5.**
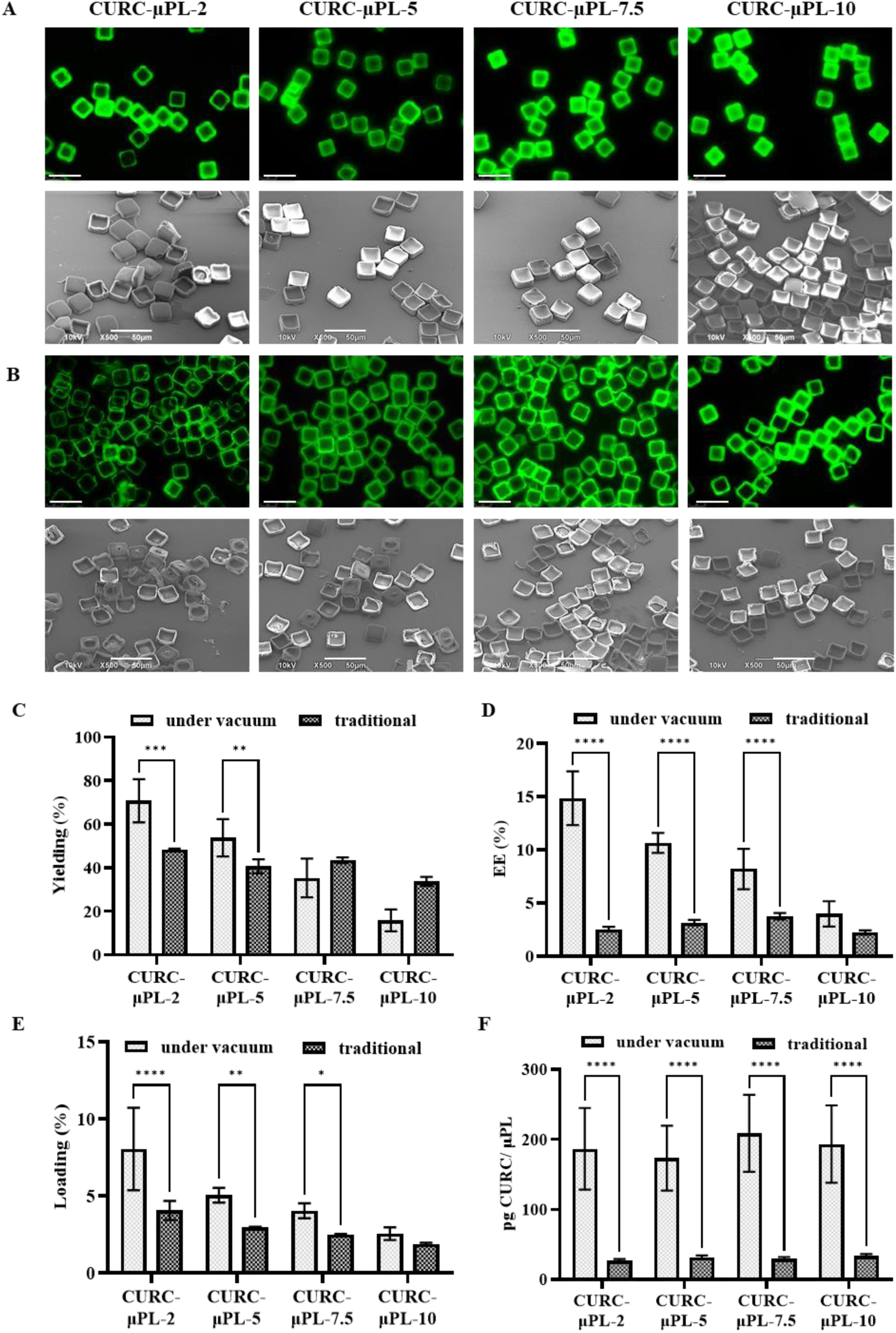
Improved morphological and pharmacological properties of CURC-μPL with vacuum and ‘cleaning’. **A.** Fluorescence microscopy and SEM (45° tilted) images of μPL obtained via the under vacuum method **B.** Fluorescence microscopy and SEM (45° tilted) images of μPL obtained via the original method; Side-by-side comparison in terms of **C.** Yielding; **D.** Encapsulation efficiency (EE); **E.** Loading; and **F.** Amount of curcumin encapsulated per µPL (pg/µPL). Results are presented as average ± SD (n = 3), ****: p <0.0001.

### Biodegradation of microPLates (μPL)

The degradation of PLGA microparticles is of fundamental importance in drug delivery and directly correlates with the release behavior ^33,34^. Diverse mechanisms, including degradation via solvent diffusion across the polymer network, diffusion through pores, swelling, hydrolysis, and erosion, may concurrently occur depending on the system composition and structure^35^. Typically, PLGA biodegradation progresses through hydrolysis of the ester bonds leading to the production of lactic-acid (LA) and glycolic-acid (GA), following a bulk erosion that can be catalyzed by the local acid environment^10^. The degradation rate is dependent on several factors ranging from the molecular weight of the polymer to the LA and GA ratios, as well as the pH of the chosen medium, the dimensions and morphology of the microparticles^36–38^. Generally, PLGA with higher molecular weight tends to exhibit a lower degradation rate, and small particles exhibit a faster degradation due to the expanded surface area which offers more sites for hydrolysis^39–43^. Even the interplay between a loaded compound (e.g. CURC) and the PLGA matrix may significantly impact polymer biodegradation profiles and release kinetics^44,45^.

Degradation studies of μPL were carried out in PBS and DI water, evaluating changes in particle morphology via SEM (**Figure 6**) and the decrease in particle number by Multisizer Coulter Counter analyses (**Figures 7** and **8**), at predetermined time points. **Figures 6A** and **6B**, respectively, show the morphological alterations that μPL underwent upon incubation in PBS and DI water as captured by SEM analysis up to 28 days. Among all the configurations investigated, CURC-µPL-2 stood out as the only one capable of maintaining a distinct square shape even after 28 days in PBS medium. This is indeed in agreement with the sustained release of CURC from CURC-µPL-2 which is expected to continue after 1 month. Differently, all the other tested configurations tend to lose their characteristic initial prismatic shape with time. Notably, the corners are smoothened and eventually the entire particle assumes a more rounded shape. As expected, the µPL degradation occurred much faster in DI water than in PBS. The hydrolysis of ester bonds between LA and GA was accelerated in water, particularly favored by its slightly acidic pH, where hydrogen ions (H^+^) enhance hydrolysis. In contrast, PBS, being a buffered solution with ions, stabilizes pH during hydrolysis, slowing down PLGA degradation^46^.

**Figure 6.**
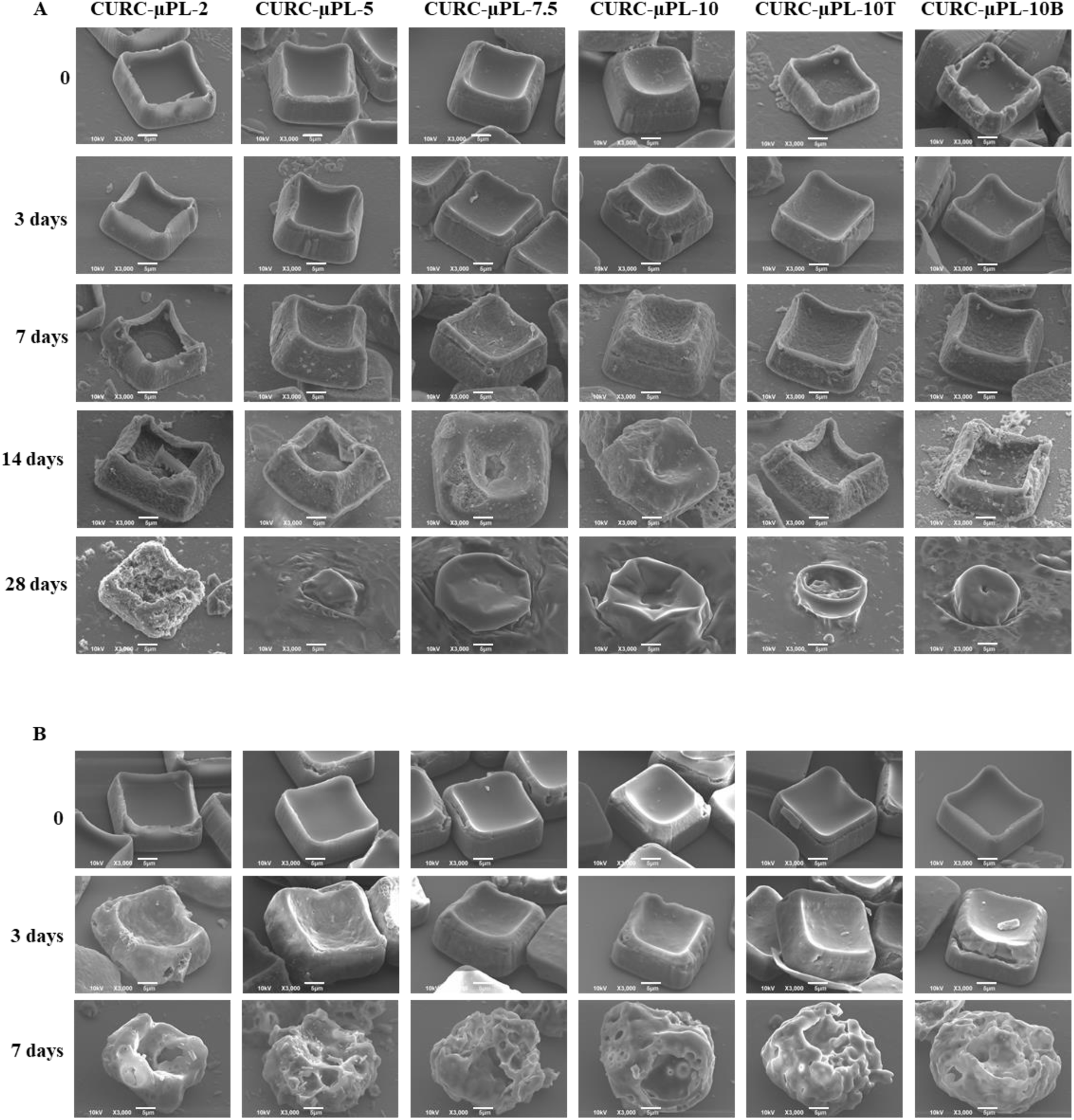
Biodegradation of curcumin loaded microPlates (CURC-μPL). A. SEM images (45° tilted) of CURC-µPL incubated in PBS (0.5 mL of PBS buffer at pH = 7.4) for up to 28 days; B. SEM images (45° tilted) of CURC-µPL incubated in deionized water (0.5 mL of DI water at pH = 5.5) for up to 7 days (scale bar: 5 μm).

**Figure 7.**
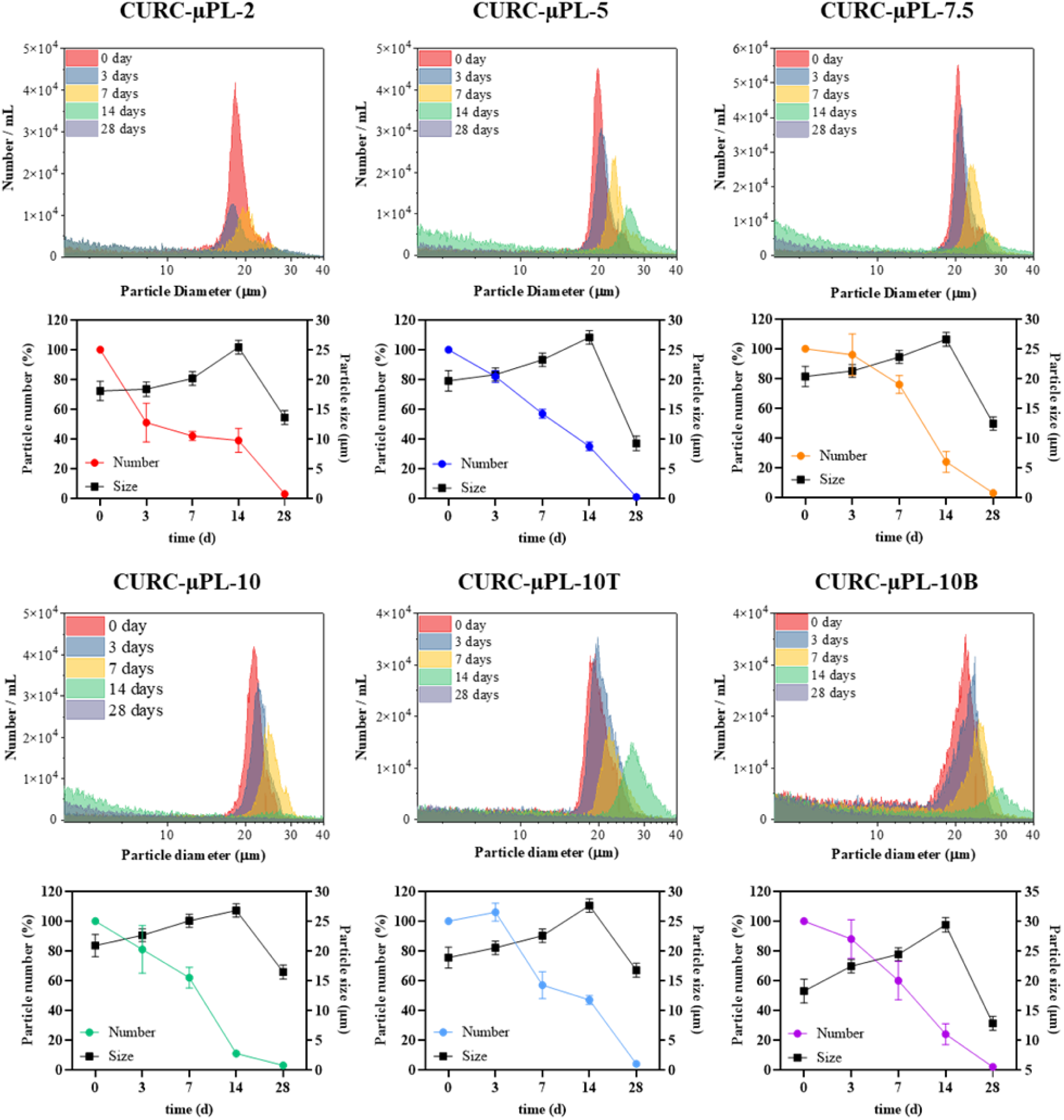
Biodegradation in PBS of six different μPL configurations. **(top)** Variation of the μPL size distribution profiles over time, via Multisizer Coulter Counter analyses; **(bottom)** Percentage of μPL number (colored line) and μPL size variation (black line) over time in PBS. Results are presented as average ± SD (n = 3).

**Figure 8.**
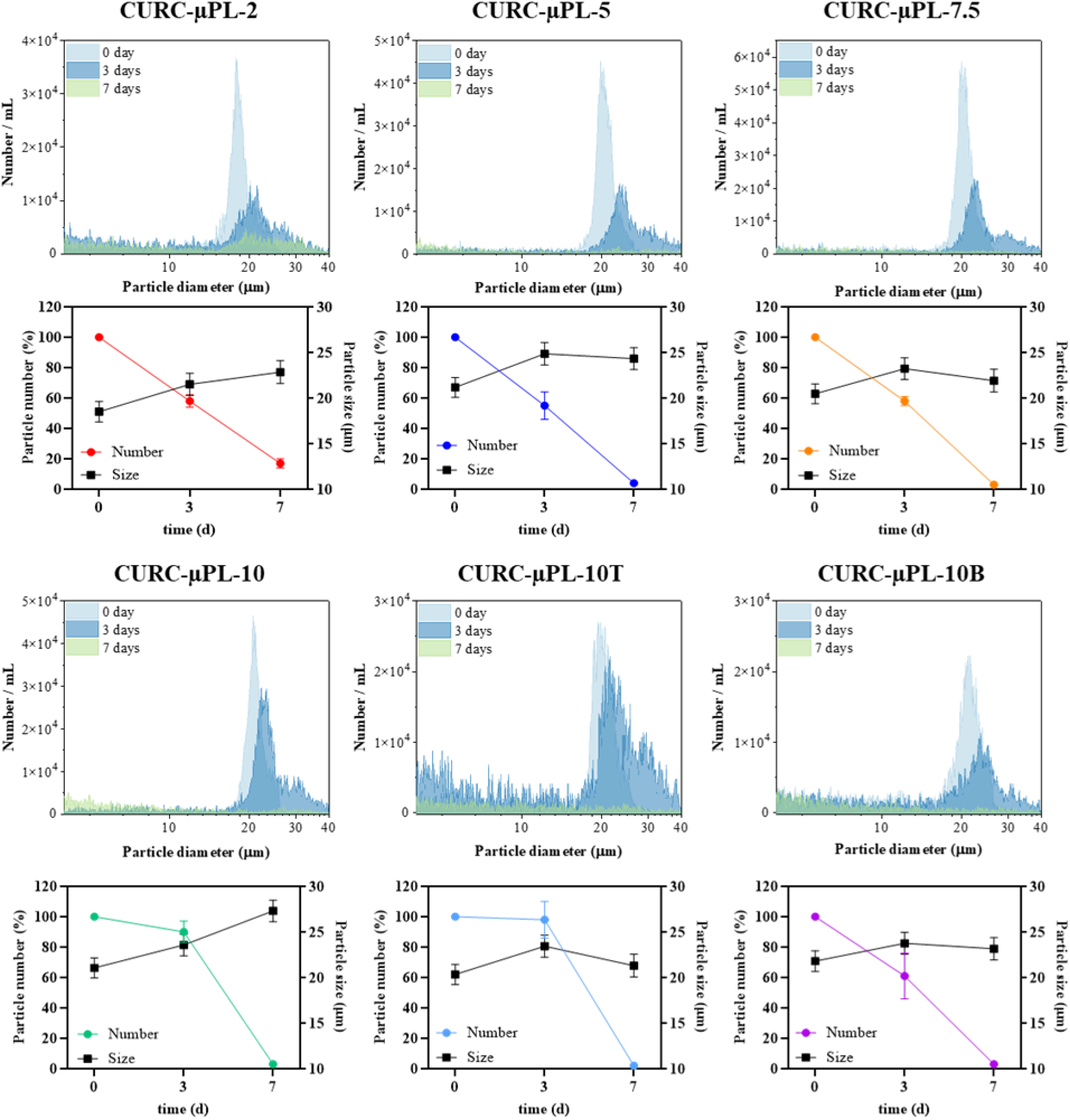
Biodegradation in DI water of six different μPL configurations. **(top)** Variation of the μPL size distribution profiles over time, via Multisizer Coulter Counter analyses; **(bottom)** Percentage of μPL number (colored line) and μPL size variation (black line) over time in DI water. Results are presented as average ± SD (n = 3).

Notably, within the same media, μPL with lower PLGA content maintained their shape longer while formulations with larger PLGA amounts released more LA and GA into the surrounding environment, decreasing local pH and favoring further PLGA degradation. This behavior is linked to the autocatalytic nature of PLGA, resulting in heterogeneous degradation due to pH gradients within the particles. Since high molecular weight PLGA follows bulk erosion, which affects the entire volume of the particle and not just the surface, the internal diffusion of acidic degradation products from the inner core to the surface and surrounding media creates a pH gradient from the center to the surface of the particles^46^. This appears to be consistent with the observations drawn from the drug release studies where μPL with larger PLGA content have faster CURC release^42,44–46^.

**Figure 7** shows the μPL size distribution (top rows) as well as the progressive reduction in μPL number and change in μPL average size over time (bottom rows) during the degradation process in PBS, as assessed via a Multisizer Coulter Counter system. For all μPL configurations, the actual characteristic size slightly grows with time up to 14 days and then rapidly decreases to values around 15 µm. This is due to the softening of the μPL matrix and water uptake. At the same time, the percentage of μPL with a well-defined square shape, as determined by the Multisizer Coulter Counter system, decreases steadily over time. CURC-µPL-2 has a quite unique behavior where a sudden reduction in the number of particles was observed up to 3 days, followed by a plateau up to 14 days and, eventually, a complete degradation after 28 days. In DI water, all µPL configurations exhibited a rapid collapse already within the first week of observation (**Figure 8**).

It is here important to note that this study highligths the critical importante of factors like polymer concentration and particle shape in modulating release kinetics. These are vital considerations for designing drug delivery systems with specific therapeutic objectives.

## Conclusions

In this study, square 20×20 μm PLGA microPLates (µPL) were produced using a top-down template-based fabrication strategy modified by the inclusion of a vacuum-chamber step for the controlled evaporation of the organic solvent. Following this approach, microparticles with different polymeric content and obtained by different purification methods were extensively characterized for their morphological, physico-chemical, and pharmacological properties. It was shown that µPL with low PLGA content were associated with higher fabrication yielding (70% for the µPL-2), higher encapsulation efficiency (∼ 15% for the CURC-µPL-2), and lower rates for drug release (∼ 60% for the CURC-µPL-2 after 10 days) and degradation. In contrast, µPL with higher PLGA content were associated with lower fabrication yielding (10% for the CURC-µPL-10), lower encapsulation efficiency (∼ 1% for the CURC-µPL-10), and higher rates for drug release (> 80% for the CURC-µPL-2 after 10 days) and degradation. Moreover, by using the Weibull and Korsmeyer-Peppas models, it was observed how CURC release from low PLGA content µPL is mostly dominated by Fickian diffusion, whereas in µPL with higher PLGA content both diffusion and polymer erosion contribute to CURC release. In summary, the proposed fabrication approach, along with the extensive characterization analyses, present the PLGA-µPL as a versatile drug delivery system where physico-chemical and pharmacological properties could be tailored to accommodate different biomedical applications.

## AUTHOR INFORMATION

### Corresponding Author

Paolo Decuzzi, PhD – Laboratory of Nanotechnology for Precision Medicine, Fondazione Istituto Italiano di Tecnologia, Via Morego 30, Genoa 16163, Italy; Division of Oncology, Department of Medicine and Department of Pathology, Stanford University School of Medicine, Stanford, CA, USA.

## Author Contributions

‡Denise Murgia and Bianca Martins Estevão contributed equally. All authors have given approval to the final version of the manuscript.

## Notes

The authors declare no competing financial interest.

## Acknowledgments

The authors acknowledge partial support from the European Union’s Horizon 2020 Research and Innovation program under the Marie Skłodowska-Curie Grant 754490 – MINDED project and Grant 872648 – MEPHOS project. The authors also recognize the support of the Italian Institute of Technology and its Facilities.

